# A unified physiological framework of transitions between seizures, sustained ictal activity and depolarization block at the single neuron level

**DOI:** 10.1101/2020.10.23.352021

**Authors:** Damien Depannemaecker, Anton Ivanov, Davide Lillo, Len Spek, Christophe Bernard, Viktor Jirsa

## Abstract

The majority of seizures recorded in humans and experimental animal models can be described by a generic phenomenological mathematical model, The Epileptor. In this model, seizure-like events (SLEs) are driven by a slow variable and occur via saddle node (SN) and homoclinic bifurcations at seizure onset and offset, respectively. Here we investigated SLEs at the single cell level using a biophysically relevant neuron model including a slow/fast system of four equations. The two equations for the slow subsystem describe ion concentration variations and the two equations of the fast subsystem delineate the electrophysiological activities of the neuron. Using extracellular K^+^ as a slow variable, we report that SLEs with SN/homoclinic bifurcations can readily occur at the single cell level when extracellular K^+^ reaches a critical value. In patients and experimental models, seizures can also evolve into sustained ictal activity (SIA) and, depolarization block (DB), activities which are also parts of the dynamic repertoire of the Epileptor. Increasing extracellular concentration of K^+^ in the model to values found during experimental status epilepticus and DB, we show that SIA and DB can also occur at the single cell level. Thus, seizures, SIA and DB, which have been first identified as network events, can exist in a unified framework of a biophysical model at the single neuron level and exhibit similar dynamics as observed in the Epileptor.

**Author Summary:** Epilepsy is a neurological disorder characterized by the occurrence of seizures. Seizures have been characterized in patients in experimental models at both macroscopic and microscopic scales using electrophysiological recordings. Experimental works allowed the establishment of a detailed taxonomy of seizures, which can be described by mathematical models. We can distinguish two main types of models. Phenomenological (generic) models have few parameters and variables and permit detailed dynamical studies often capturing a majority of activities observed in experimental conditions. But they also have abstract parameters, making biological interpretation difficult. Biophysical models, on the other hand, use a large number of variables and parameters due to the complexity of the biological systems they represent. Because of the multiplicity of solutions, it is difficult to extract general dynamical rules. In the present work, we integrate both approaches and reduce a detailed biophysical model to sufficiently low-dimensional equations, and thus maintaining the advantages of a generic model. We propose, at the single cell level, a unified framework of different pathological activities that are seizures, depolarization block, and sustained ictal activity.

## Introduction

Seizures are part of the repertoire of built-in activities of neuronal networks as they can be triggered in most brain regions from most species (Jirsa et al. 2014). Several conceptual frameworks have been proposed to explain seizure dynamics (Naze 2015; Naze et al. 2015; Soltesz and Staley 2008; Staley 2015; Stefanescu et al. 2012; Y. Wang et al. 2017; Wendling et al. 2016). The predominant framework assumes that the majority of seizure onsets and offsets correspond to bifurcations in the electrophysical variables (Jirsa et al. 2014; Saggio et al. 2017), although there exist other non-bifurcation types (Blenkinsop et al. 2012). This framework has been generalized by Saggio and colleagues (Saggio et al. 2017, 2020). A phenomenological mathematical model, called The Epileptor, describes the dynamics of a majority of seizures recorded in drug-resistant patients, and most seizures recorded in experimental models (Jirsa et al. 2014; Saggio et al. 2020). A qualitative analysis of the Epileptor reveals that seizures, SIA and DB co-exist, and that multiple types of transitions from one type of activity to the other are possible, as verified experimentally (El Houssaini et al. 2015; Houssaini et al. 2020; Saggio et al. 2017). Since it is phenomenological, the Epileptor model does not provide direct insight regarding underlying biophysical mechanisms. The phenomenological model imposes strong constrains in terms of dynamics. Numerous neuronal network models, including biophysically realistic ones, have been developed to study seizures, SIA or DB mechanisms (Holt and Netoff 2013; V.K. Jirsa et al. 2017; Kim and Nykamp 2017; Krishnan et al. 2015; Lytton 2008; Naze 2015; Proix et al. 2014; Stefanescu et al. 2012; Traub and Wong 1982). These models contain too many parameters and variables to perform a detailed bifurcation analysis, thus preventing bridging the gap between phenomenological and biophysical approaches. However, with the guidance of phenomenological modeling, design of neuronal spiking network including several biophysical features has been performed (Naze et al. 2015). In this work, transitions between states of the neuronal network are ensured by slow variable representing extracellular environmental fluctuation.

Although seizures and DB are generally observed at the neuronal network scale, their dynamical equivalence can be found at the single cell level (Bikson et al. 2003; Bragin et al. 1997; Chizhov et al. 2018; Cressman et al. 2009; Hübel and Dahlem 2014; Kager et al. 2000; Lietsche et al. 2016; McCormick and Contreras 2001), which can be used to study human epilepsy (Merricks et al. 2015; Tankus 2016; Truccolo et al. 2011). Moreover, previous works (Viktor K. Jirsa and Stefanescu 2011; Montbrió et al. 2015; Naze et al. 2015) show that dynamical features are preserved when going from the weakly coupled network to the single cell level. It therefore seems appropriate to consider a biophysical model at the single cell scale exhibiting dynamic properties identified in the generic model and in which the transitions between the different states are provided by a slow variable describing the variations within the extracellular milieu. Bursting activity in neurons can be described in terms of bifurcations (E. Izhikevich 2007; E. M. Izhikevich 2000), and different single cell biophysical models have been proposed, which can model SLE and DB, but not SIA (Barreto and Cressman 2011; Chizhov et al. 2018; Cressman et al. 2009; Hübel and Dahlem 2014; Kager et al. 2000; Øyehaug et al. 2012; Ullah and Schiff 2010; Y Wei et al. 2014; Yina Wei, Ullah, Ingram, et al. 2014), although, to the best of our knowledge, these activities have not been observed experimentally in isolated neurons. They are slow/fast systems, where a slow subsystem drives the fast subsystem between different states. In such models, the studied fast subsystem delineates the neuronal membrane electrophysiological activities. The slow subsystem can be represented by variations of different slow variables including ion concentration (Barreto and Cressman 2011; Chizhov et al. 2018; Cressman et al. 2009; Hübel and Dahlem 2014; Kager et al. 2000; Øyehaug et al. 2012; Y Wei et al. 2014; Yina Wei, Ullah, Ingram, et al. 2014), oxygen level (Yina Wei, Ullah, and Schiff 2014; Yina Wei, Ullah, Ingram, et al. 2014), volume (Øyehaug et al. 2012; Y Wei et al. 2014) and interaction with glial cells (Hübel and Dahlem 2014; Øyehaug et al. 2012). These models provide mechanistic insights, in particular how the slow variable influences neuronal activity, including the transitions from “healthy” regimes to “pathological” ones like SLEs and DB. However, none of these models, including the extracellular slow variations, could be reduced to four variables, while presenting a bursting pattern corresponding to the most common ones encountered at the network level, i.e. the SN/homoclinic pair (Jirsa et al. 2014). The reduction of a model to a small number of dimensions while maintaining the link with physical quantities, makes it possible to establish a direct link between biophysics and a simple dynamical description. Since SLEs, SIA and DB pertain to the dynamic repertoire of The Epileptor and of biological neuronal networks (El Houssaini et al. 2015; Saggio et al. 2017), biophysical models should be able to reproduce all three types of activities. The goal of the present study is to identify candidate mechanisms from physiology robustly leading to the time scale separation and trajectories in SLEs. We will take guidance from the careful dynamic analyses performed in previous works.

In generic models, the dynamics is well understood (Jirsa et al. 2014; Saggio et al. 2017) but these models rarely offer direct biophysical insight as they use abstract parameters. Important works have been done to understand the link between phenomenological and biophysical models (Bernard et al. 2014; Naze et al. 2015). In order to explore the dynamics repertoire, the high dimensionality of detailed biophysical models must be reduced. A minimal model of interictal and SLEs has been introduced as Epileptor-2, using increase in [K]_o_ to trigger burst discharges and restoration of the K^+^ gradient via the sodium-potassium pump to stop SLEs (Chizhov et al. 2018). The work of Saggio et al. (Saggio et al. 2017) is a generalization of the dynamics found in Epileptor and Epileptor-2. However, Epileptor-2 does not produce the SN/homoclinic bifurcation consistently found experimentally, and the model does not generate SIA or DB. Moreover, when the fast discharges stop, the slow variables of Epileptor-2 continue oscillating, whereas that of Epileptor 1 do not. This can be understood on the basis of the results of Saggio et al.(Saggio et al. 2017). In the present reduction, we used a Hodgkin-Huxley-like single cell model, and we imposed several constraints: SLE, SIA and DB (El Houssaini et al. 2015), as well as the SN/homoclinic bifurcation must be present. We have thus generated a reduced slow/fast system preserving biophysical representation and satisfying these constraints. A variable acting on a slow time scale is necessary to drive the system through different activities (e.g. from SLE to DB). In a cell, numerous processes can occur on a slow time scale and act as a slow variable, including changes in ion concentration, metabolism, phosphorylation levels, and transcription. Ion homeostasis regulation is critical to maintaining neuron function, and many sub-cellular mechanisms are involved in this process (Dubyak 2004). Fluctuations of ion concentrations in the extracellular space modulate the electrophysiological activity of a single neuron (Cressman et al. 2009; Yina Wei, Ullah, and Schiff 2014). The present work focuses on extracellular potassium ([K]_o_) concentration because it increases during seizures (de Curtis et al. 2018; Fisher et al. 1976; Fröhlich et al. 2008; Lux et al. 1986; L. Wang et al. 2016), even in the absence of synaptic activities (de Almeida et al. 2008; Jefferys and Haas 1982). Augmentation of [K]_o_ is also observed in head injury (Katayama et al. 1990; Reinert et al. 2000), which can be a starting point for epilepsy (Lowenstein 2009). Computational simulations show that potassium could be responsible for local synchronization (Durand et al. 2010) and is an important parameter in neural dynamics (Barreto and Cressman 2011; Cressman et al. 2009; Ullah and Schiff 2010; Yina Wei, Ullah, and Schiff 2014). In addition, in experimental models, the transition to DB correlates with a much larger increase of [K]_o_ as compared to SLEs (El Houssaini et al. 2015; Gloveli et al. 1995). We here consider the slow modulatory effects of [K]_o_ variations. In our model, the slow sub-system describes ionic concentration variations. The fast subsystem characterizes the dynamics of trans-membrane ion flows through voltage-gated and the sodium-potassium pump, and so allows tracing the membrane potential. We report that this single cell model accounts for the SN/homoclinic bifurcation pair and that it reproduces SLEs, SIA and DB, reproducing patterns found in single neurons recorded experimentally during seizures.

## Results

Our goal was to construct a biophysical model at single neuron level that can reproduce the different firing patterns recorded when extracellular potassium is experimentally increased, while keeping it sufficiently simple to allow a bifurcation analysis. The model is schematized in Fig.1 (see methods section for the equations). It is a simplification of the classical Hodgkin-Huxley formalism, which also includes close surrounding environment thanks to a description of three compartments (external bath, extracellular space, intracellular space).

**Figure 1:**
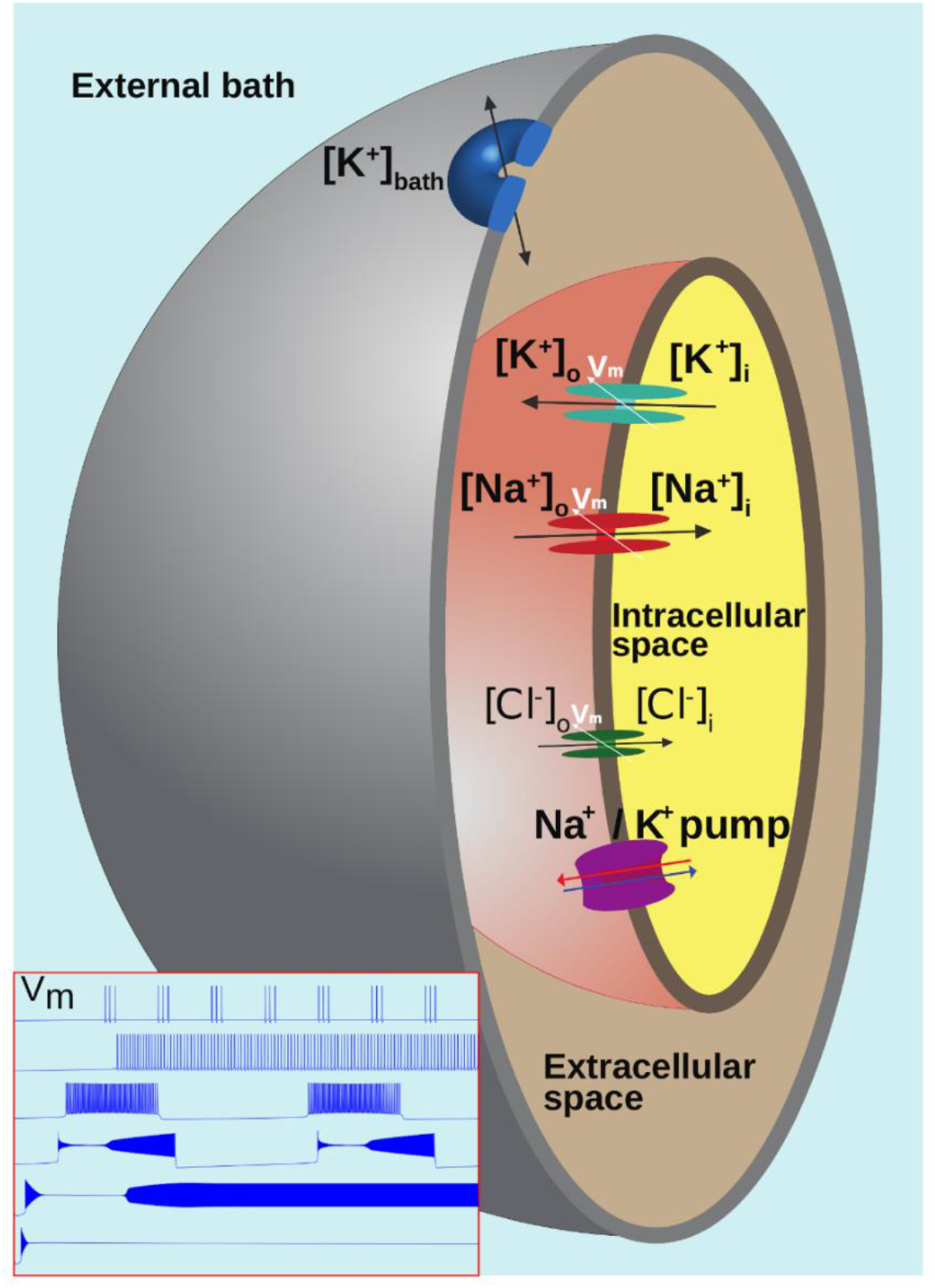
Diagram of characteristics and mechanisms described by the model. Three compartments are represented. A passive diffusion of potassium exists between the external bath and the extracellular space. Na^+^, K^+^ and Cl^-^ ions can be exchanged between the extracellular and intracellular compartments via the Na/K-pump and voltage-gated channels. This model can reproduce the typical patterns of the membrane potential V_m_, shown in the bottom left subplot, including tonic firing, bursting, seizure like events (SLE), sustained ictal activity (SIA) and depolarization block (DB).

Numerous experiments show that seizures and SLEs are associated with an increase in [K]_o_ (Fisher et al. 1976; Fröhlich et al. 2008) and that increasing of external [K] can trigger SLEs (S. F. Traynelis and Dingledine 1988; Stephen F. Traynelis and Dingledine 1989). The model presented here takes into account the regulation of potassium, via the possible diffusion towards the external bath compartment and its associated potassium concentration [K]_bath_. Changing [K]_bath_ parameter will strongly influence the regulation of extracellular potassium by allowing or not the removal of excess potassium from the extracellular compartment. When [K]_bath_ is low, the bath compartment can pump out the extracellular potassium; but it fails to do so when it is saturated by potassium. We thus explored the response of the model as the concentration of [K]_bath_ was increased. The gradual increase in potassium led to 7 sequential qualitative firing patterns: Resting State (RS), Spike Train (ST), Tonic Spiking (TS), Bursting, Seizure-like events (SLE), Sustained Ictal Activity (SIA), and Depolarization Block (DB) (Note that what is called here Spike Train also corresponds to another type of burster from a dynamical point of view (E. Izhikevich 2007; Saggio et al. 2017)). The corresponding changes of membrane potential for all these patterns are shown in Fig.2.

**Figure 2:**
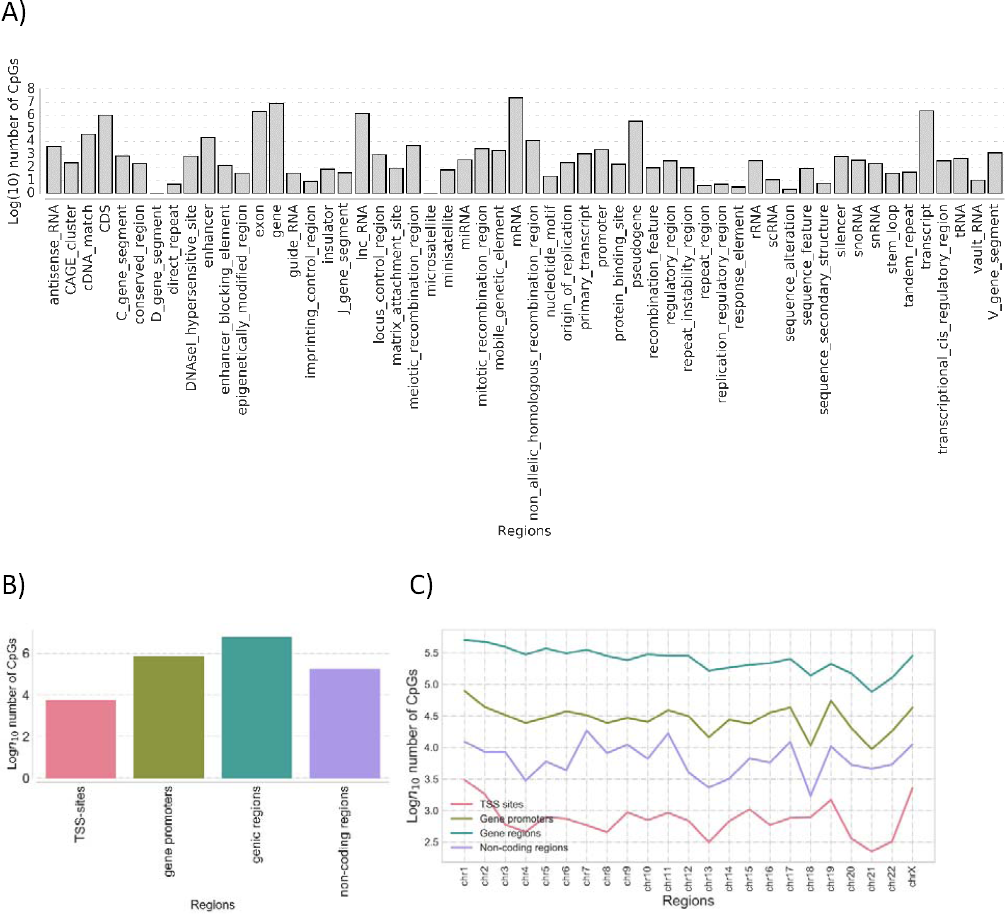
qualitative mode of behavior of the membrane potential and Nernst potentials. In blue: time series of the membrane potential V_m_ for the following patterns of activity: (a) Spike train, (b) Tonic spiking (TS), (c) Bursting, (d) Seizure-like event (SLE), (e) sustained ictal activity, (f) Depolarization Block (DB). In red: Nernst potential of sodium, in green: Nernst potential of potassium with specific Y axis on the right side. If the value of [K]_bath_ stays below 6 mM, the system remains in resting state around -72 mV. Specific patterns of activities start to appear with a diminution of the Nernst potential of sodium and an increase of the Nernst potential of potassium. When periodic events are occurring (panels c and d), oscillations can also be observed in the Nernst potential of both ions.

The number of firing patterns is higher than in the original Hodgkin-Huxley model. This is due to the fact that the model takes into account the variations of concentration, as evidenced by the variation of the Nernst potentials. The changes in Nernst potentials for sodium and potassium ion species are shown in Fig.2. The simulations are initialized with values observed in a “healthy” resting situation, it is therefore possible to observe the transition linked to the increase in the parameter [K]_bath_. In some cases, the Nernst potentials display a transient change before reaching a sustain low amplitude oscillations following action potentials, as observed during RS, TS, SIA and DB. During periodic events, (ST, Bursting, SLE), larger oscillations are observed in Nernst potentials. These oscillations are directly linked to the observed oscillations in the slow variables of the model (Eq.(3) and Eq.(4)) describing concentration changes. The rate of oscillation of the slow variables thus explain the duration of periodic events, in line with the assumed essential role of ionic homeostatic regulation.

Each of the firing patterns can be associated to a different behavior, observable experimentally at different scales. The correspondence is established on the basis of their shape and their order of appearance as [K]_bath_ is increased. Tonic and bursting patterns are prototypical. We consider the activity shown on Fig. 2d as SLE at the neuronal scale, as it is similar to the activity typically recorded in individual neurons (Haglund and Schwartzkroin 1990), in particular the transient episode of depolarization block, in different experimental preparations during SLEs at the network scale (e.g. Fig. 6 in (Uva et al. 2013); Fig. 1 in (Bikson et al. 2003) or Fig. 8 in (Jirsa et al. 2014)). Although it is possible to generate SIA *in vitro* (Quilichini et al. 2002), the corresponding intracellular activity is not known, however the sustained firing pattern in the model cell resembles the regular field activity recorded during status epilepticus like events *in vitro* (Quilichini et al. 2002). The sustained DB at the single cell level corresponds to what is observed experimentally during network spreading depolarization when [K]_o_ reaches high levels (G G Somjen 2001).

Increasing [K]_bath_ leads to different regimes of variation of external potassium (Fig.3). These different regimes are associated with a specific dynamic (i.e. type of bifurcation) of the excitability of the membrane. It is therefore possible to link the membrane potential to the variations in extracellular potassium, because of exchanges existing between compartments (i.e. via the slow variable), as shown in Fig.4. In the next subsection, we detailed these dynamical interactions for the different patterns of activity, following the order of appearance when [K]_bath_ increase.

**Figure 3:**
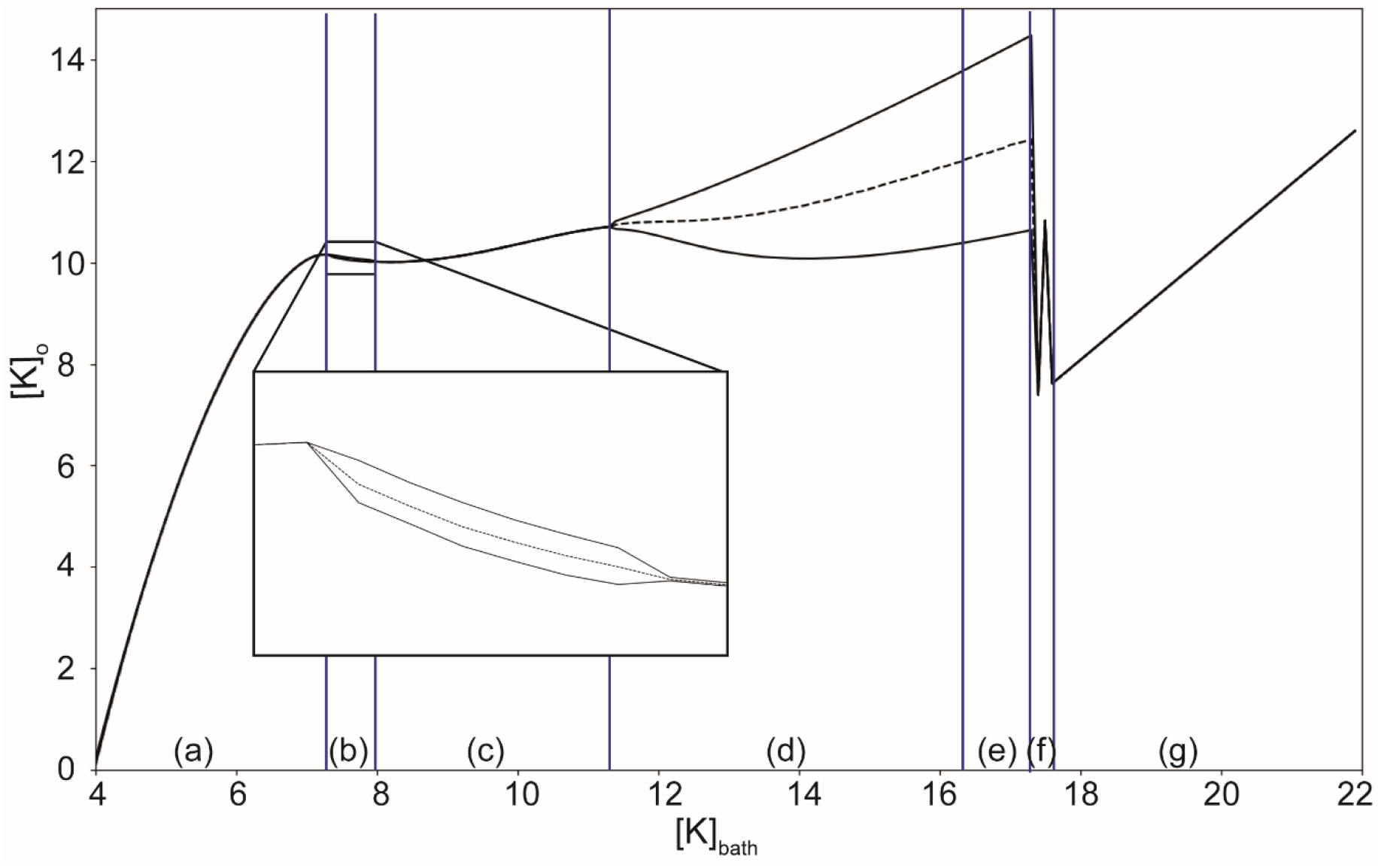
Variation of extracellular potassium concentration as a function of [K]_bath_. Minimal and maximal external potassium [K]_o_ and mean (dash line) concentration observed during simulations done for different values of the parameter [K]_bath_. Due to diffusion from the external bath, increasing [K]_bath_ leads to variations in [K]_o_. Different patterns are observed for each range of [K]_bath_: (a) resting state, (b) spike train, (c) regular spiking, (d) burst, (e) seizure like event, (f) sustained ictal activity, (g) depolarization block. The periodic events (spike train, burst and seizure-like event) correspond to the range of [K]_bath_ where [K]_o_ periodically oscillates.

**Figure 4:**
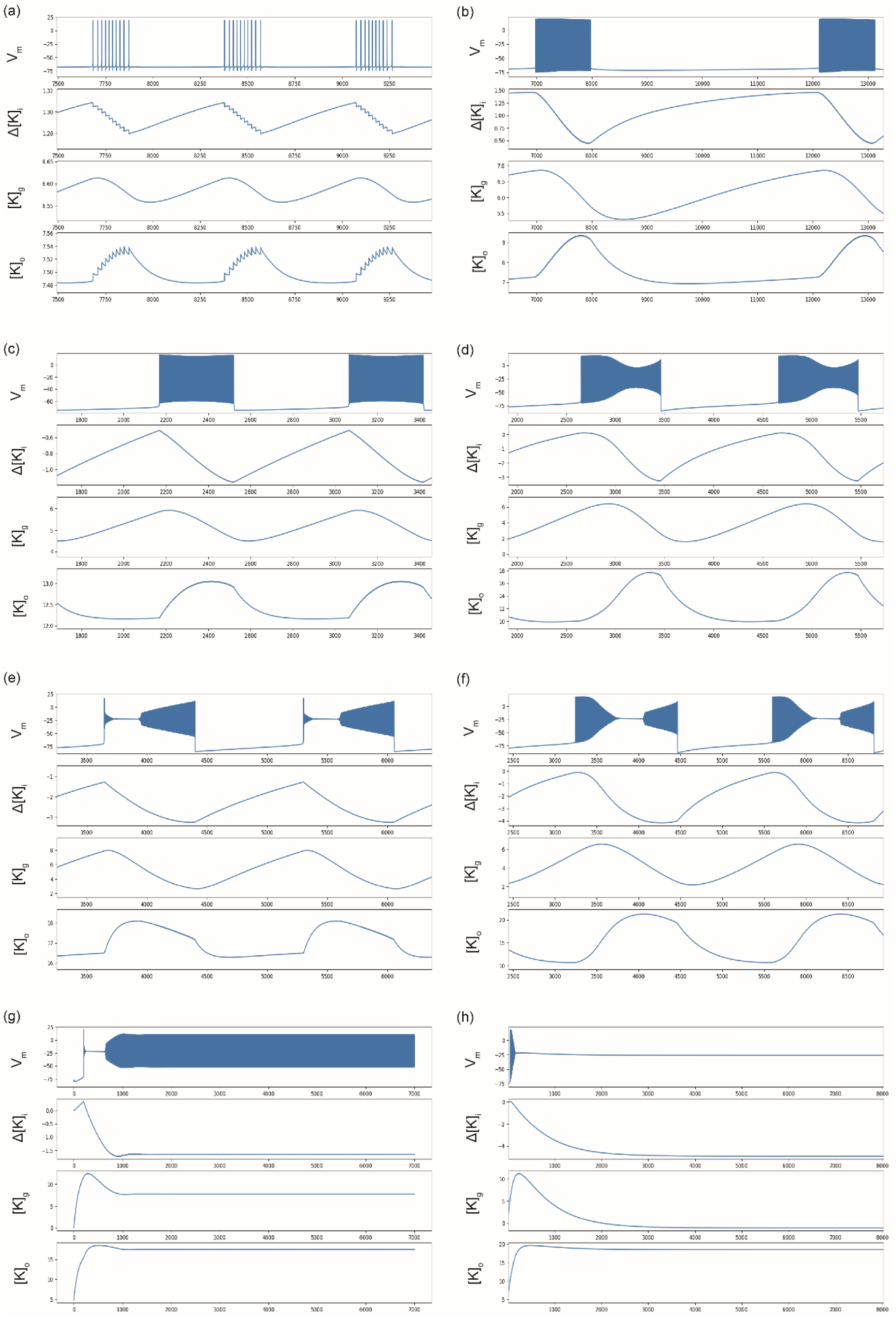
Time series of membrane potential, Δ[K]_i_, [K]_g_, and [K]_o_. obtained thanks to numerical methods using SciPy library (Millman and Aivazis 2011), x-axis in millisecond. (a) spike train with [K]_bath_ = 7.5 mM, (b) spike train with [K]_bath_ = 7.5 mM and γ = 0.04, ε= 0.002, (c) Burst with [K]_bath_ = 12.5 mM, (d) Burst with [K]_bath_= 12.5 mM, and γ = 0.06, ε = 0.002, (e) SLE with [K]_bath_ = 16 mM, (f) SLE with [K]_bath_ = 16 mM and γ = 0.08, ε = 0.0008, (g) SIA with [K]_bath_= 17.5 mM, (h) DB with [K]_bath_ = 20 mM. If not specified, the parameter values used here are the reference parameters described in the method section. Variations of Δ[K]_I_ and [K]_g_ induce different patterns of activity. The combined effects lead to the observed variations in [K]_o_. The time scale of the slow variables γ and ε influence the shape of V_m_ allowing the system to exhibit SN or SNIC bifurcation at the onset of the events.

### Resting states, spike train and tonic spiking in low [K]_bath_

Resting state is found when [K]_bath_ is around the normal value of [K]_o_ (called [K]_0,o_, see Methods section). If [K]_bath_ is smaller than [K]_0,o_ the membrane potential slowly hyperpolarizes, due to a diffusion of potassium in the direction of the external bath. When [K]_bath_ slightly increases (> 7 mM), ST appears through a SNIC bifurcation. The offset is also a SNIC bifurcation. In this case, the onset and offset bifurcations can be easily identified by their characteristic features (Saggio et al. 2017), and confirmed by numerical methods (using SymPy (Meurer et al. 2017) and SciPy (Millman and Aivazis 2011) libraries). With higher value of [K]_bath_ (> 8 mM), TS occurs. In this condition, [K]_o_ stabilizes (Fig.3), and the neuron fires at a constant frequency (Fig. 2b). The occurrence of regular spiking due to an increase of [K]_o_ through diffusion from the bath is consistent with experiments [29].

### Bursting and Seizure-like events

Bi-stable behavior occurs when the slow system starts to oscillate when [K]_bath_ is further increased. The model (with parameters listed in Table 1) displays bursting and SLE, successively. Bursts are square-wave bursts (SN/Homoclinic bifurcations) and SLEs also show SN and Homoclinic bifurcations at onset and offset, respectively (See supplementary information: S1, S2, S3, S4). Here, the slow subsystem oscillates in a self-sustained manner (Fig. 4a-f), generating recurrent bursting or SLEs, with important variations of [K]_o_, due to oscillations in the slow subsystem. The combined effects of oscillations of Δ[K]_I_ and [K]_g_ explain the changes in the Nernst potential of potassium (and sodium, which is linked to potassium in the model), thus changing neural excitability. During spiking activity, voltage-gated potassium channels open increasing potassium current I_K_. The influence of I_K_ in the equation of Δ[K]_I_ (eq.3), explains the decrease of Δ[K]_I_, hence the increase of [K]_o_ through equations (eq.16) and (eq.20). This is consistent with the observations described in (Fisher et al. 1976). The increase in [K]_o_ starts with the occurrence of burst and SLEs. Thus, it is not the cause of the event but a consequence of homeostasis dysregulation (i.e. augmentation of [K]_bath_).

**Table 1.**
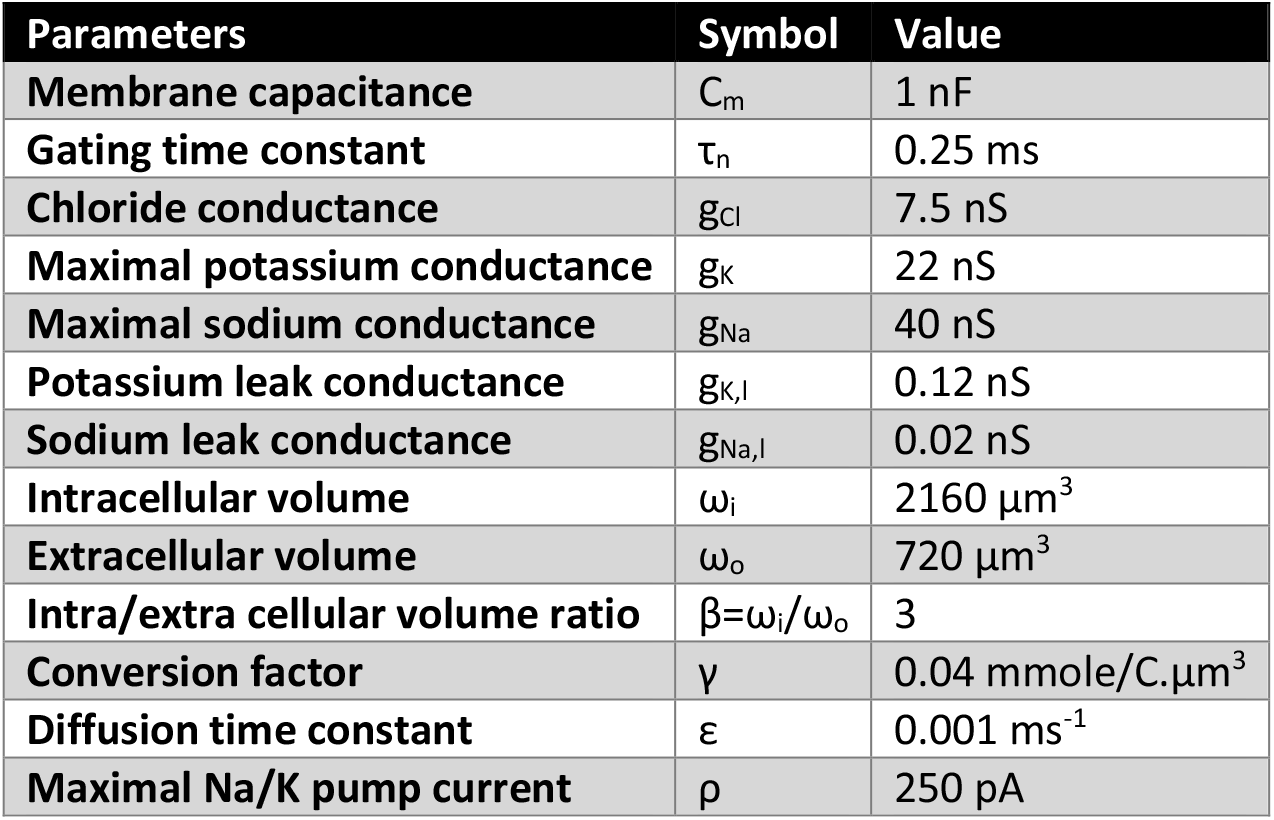
Parameters values.

### Steady states, SIA events and DB, in high [K]_bath_ conditions

SIA events (Fig. 2g) appears for [K]_bath_ around 17.5 mM (Figs. 3 & 4), i.e. above the threshold value for SLEs as reported experimentally (El Houssaini et al. 2015). If no other mechanisms act to stop it, these oscillations remain constant (analogous to refractory status epilepticus). Permanent DB occurs for even higher values of [K]_bath_ (> 18.0 mM, Fig. 3) as also reported experimentally (El Houssaini et al. 2015). In these cases, after a peak value (Fig. 4h), [K]_o_ stabilizes, explaining the short range of variation (Fig. 3). These steady-states start like a SLEs (Fig. 4e-f), then the slow variables stabilize and [K]_o_ remains constant at a high value (Figs. 4g-h).

We conclude that the model behaves as expected from the biological observations, when experimentally increasing of [K]_bath_. These simulations were obtained when using a “healthy” situation, i.e. as if recording a “control” neuron. In the next section, we model a “pathological” situation for which where the regulatory mechanisms of neuronal homeostasis are affected.

### Influence of other parameters

We then aimed to identify relevant parameters that could describe “healthy” and “pathological” states. Experimental data show that impairment in potassium buffering by glial cells leads to pathological behavior (Coulter and Steinhäuser 2015; Hubbard and Binder n.d.; Rangroo Thrane et al. 2013; Scholl et al. 2009). Three model parameters correspond to homeostasis regulation, involving two mechanisms: the ion exchange capacity between compartments (ε and γ parameters in the model), and the maximum capacity of the Na/K pump (ρ in the model). A variation of ε corresponds to a degradation of the interaction with glial cells (Coulter and Steinhäuser 2015; Kofuji and Newman 2004; Olsen et al. 2015; Walz 2000) which normally ensures the regulation of the extracellular concentration of K^+^. Homeostasis of intracellular ions is also critical, and a variation of γ corresponds to an impairment of such mechanisms, not detailed in this model, such as co-transporters and exchangers (Hille 2001; Kandel, Eric R., Schwartz James H., Jessell Thomas M. n.d.). Changes in both parameters can also be considered. In the model, they are equivalent to a change in volume while keeping the β ratio constant. Varying the time constants of the slow subsystem (ε and γ), leads to different bi-stable behaviors. Two examples are shown in Fig. 4, (b) with γ = 0.04, ε= 0.002, s (d) γ = 0.06, ε = 0.002, and (f) γ = 0.08, ε = 0.0008, in these situations the shape of the oscillations of potassium concentration are affected leading to a change in the duration of the events. For burst and SLE shown in Fig4. d and f, the model exhibits a different class of onset bifurcation. For both, a saddle-node on invariant cycle (SNIC) bifurcation at the onset and homoclinic bifurcation at the offset can be identified, thanks to their specific dynamics and resulting shapes (E. Izhikevich 2007; Saggio et al. 2017).

The other key parameter to consider is the pump rate ρ. The Na/K-ATPase is described by Eq. (8) in the model. In a biological neuron, the pump depends on ATP and during status epilepticus, the ATP concentration augments due to high needs and then decreases (Lietsche et al. 2016). The ATP concentration is not taken into account in the model, but the maximal Na/K-pump rate is modulated by the parameter ρ. This parameter also influences the shape of I_pump_ response as a function of [Na]_i_ and [K]_o_ (Fig.5a). For large values of ρ, the pump is activated for lower value of [Na]_i_ and [K]_o_ (Fig. 5a). We find that burst duration changes with ρ for a fixed [K]_bath_ (Fig. 5b), where a faster activation (higher ρ) leads to shorter bursts. The augmentation of ρ does not necessary lead to an increase of I_pump_; it affects the general dynamics of the whole system (Fig. 5c).

**Figure 5:**
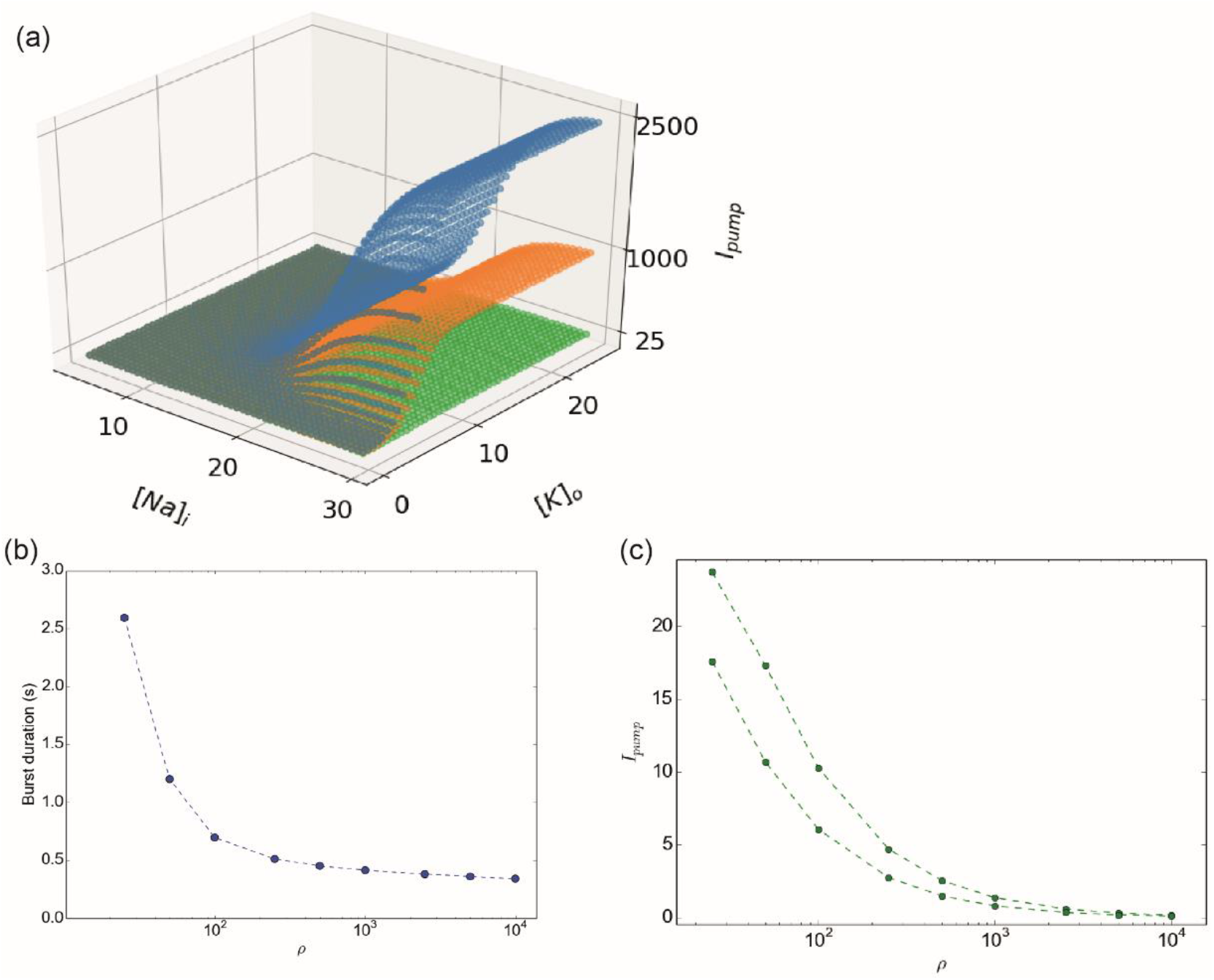
Influence of the activity of the Na/K-pump. (a) I_pump_ function for ρ= 25 (green), ρ= 1000 (orange), ρ= 2500 (blue). The initial slope when the system moves away from the concentrations at rest is affected, explaining the modification of the influence of I_pump_ in the dynamic of the system. (b) Burst duration as a function of ρ for [K]_bath_ = 14.0 mM. Bursts have shorter durations for higher value of ρ. (c) Minimal and maximal pump current, I_pump_, observed during simulation done with [K]_bath_ = 14.0 mM. The range of I_pump_ decreases for higher ρ values.

All these observations show that the model presents a behavior consistent with experimental observations. Importantly, the biophysical model is able to reproduce general patterns of activities (i.e. periodic events) as generated by the phenomenological model (El Houssaini et al. 2015). Phenomenological models, which present a minimal number of variables and parameters, allow an exhaustive study of the dynamics. The biophysical model used here contains too many parameters for an exhaustive study of the dynamics, but reducing the number of variables will allow a comparison with the generic model. In the next section, we analyze the dynamics of the model.

### Dynamical observations

The model can be divided into the fast (V, n, respectively Eq.1 and Eq.2) and the slow subsystems (Δ[K]_i_, [K]_g,_ respectively Eq.3 and Eq.3). The slow system can oscillate and drive the fast system between different behaviors, in particular switching between resting state and fast oscillations to obtain bursting-like activity. This type of phenomenon, corresponding to slow fast systems, has been extensively studied from a theoretical point of view, in particular for neural activity (E. Izhikevich 2007; Saggio et al. 2017). In this subsection, to allow a better correspondence with the theoretical framework, we call burster a system allowing these periodic events. To create the oscillation in the slow subsystem, theoretical works show that two mechanisms are possible (E. Izhikevich 2007; Saggio et al. 2017): Slow-Wave (SW) burster, where the slow subsystem is made of two equations, independent of the fast system, or Hysteresis-Loop (HL) burster where the slow subsystem is made of only one equation that depends on the fast system. Each has typical onset/offset bifurcation pairs. These specific paths for bursting have been identified in the generic model (Saggio et al. 2017), and are reproduced in fig.6a. We first verified if the relations between the equations of the slow and fast systems allow the existence of the mechanisms described previously. In our model, two equations describe the slow subsystem (Eq.(3), (4)). Because I_K_ (Eq.(6)) depends on V and n, the Eq.(3), depends on the fast system. This corresponds to a relation that exists in an HL burster. The second equation of the slow subsystem, Eq.(4), also depends on the Eq.(3), through the Eq. (20). Thus, there exists a relation between the two equations of the slow system, enabling oscillation such as in a SW burster. These relations between the variables of our model allow obtaining the two types of bursters previously described.

**Figure 6:**
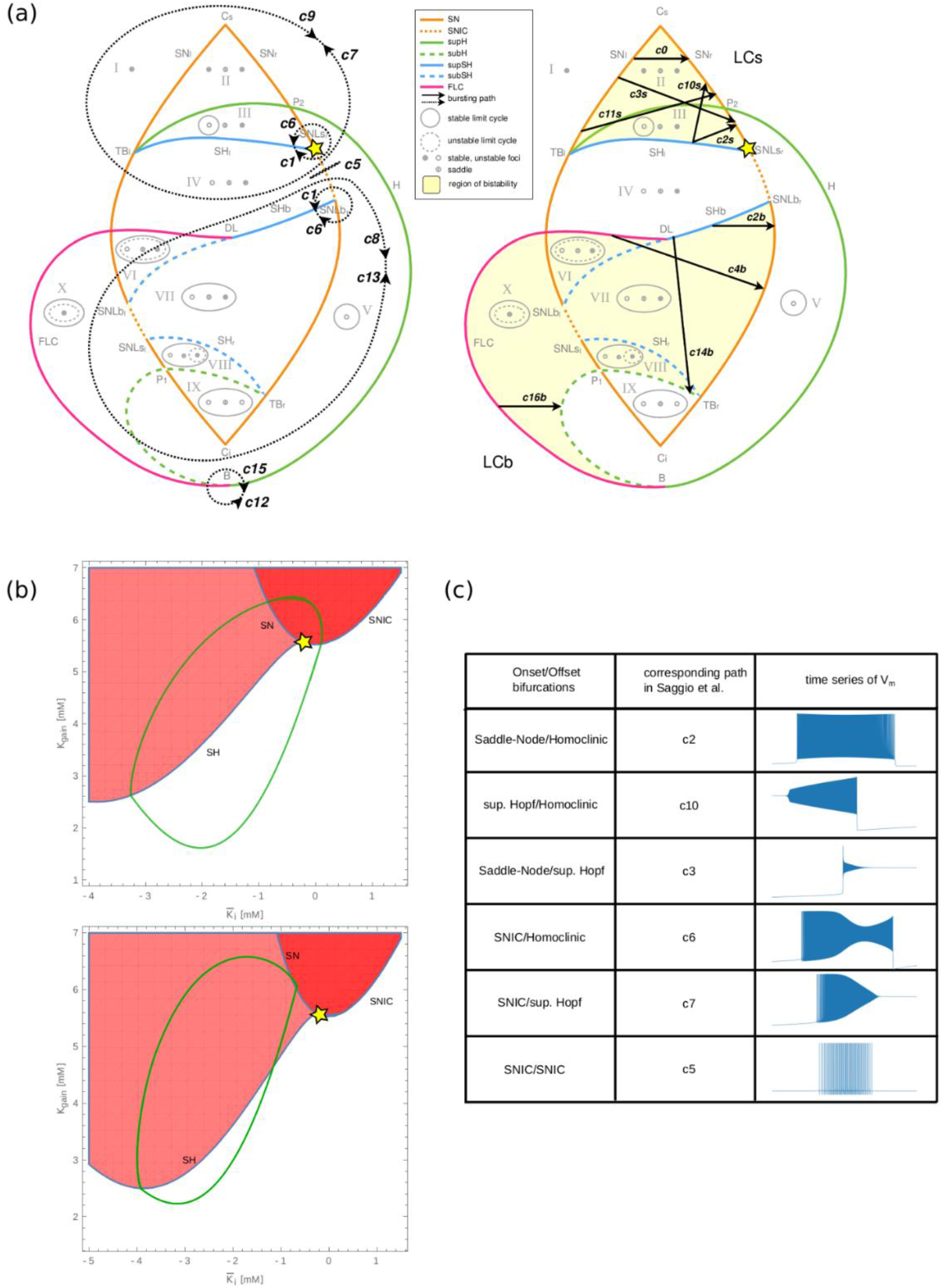
Comparison with the generic model. (a) Paths for bursting activity of the generic model proposed by Saggio et al. adapted with authorization from (Saggio et al. 2017), for hysteresis-loop burster (left) and slow-wave burster (right), the yellow star corresponds to the center of the region captured by our model. (b) Bifurcation diagram of our model, where the white area corresponds to ‘resting state only’ region, the dark red corresponds to a depolarized region, and the light-red region is the region of bi-stability. The yellow star corresponds to the point also found in the generic model, where the SH, SNIC and SN bifurcations intersect. In the top diagram, the green line corresponds to the path taken by the burster, in the bottom one to the path taken by the SLE. (c) Classes of bursters found in the model, and the corresponding path in the generic model.

We therefore tested for possible correspondences between our model and the generic model. We were able to identify the regions in the generic model capturing the dynamics reproduced by our model in Fig.6a. The center of the region of interest has been marked with a yellow star in fig. 6a. for the generic model and its correspondence in the bifurcation diagram of our model in Fig. 6b. In this bifurcation diagram we show two possible paths of our model, for burst behavior (Fig. 6b, top) and for SLE (Fig. 6b, bottom). It crosses regions of stable resting state (in white), depolarized (red), and bistable (light red). It is therefore possible to establish a non-exhaustive list of the correspondences between the paths of the two models. The paths for the periodic events have been listed in Fig. 6c. The spike train, Bursting and SLE behaviors correspond to paths, c5, c2 and c10, respectively. The bursting behavior with changes in ε and γ (Fig. 4b) that represents the SNIC/SH bifurcation corresponds to the path c6. The model proposed here, consistent with biophysics, fits into the framework of the generic model.

Since our biophysical model reproduces the bifurcations of the generic model for different types of network activities, it becomes possible to investigate the ionic mechanisms underlying the onset/offset bifurcations. The fast subsystem can be described fixing all parameters (tables 1 and 2) and considering the two slow variables as parameters. Fixed points can thus be found for different values of Δ[K]_i_ and [K]_g_ as shown in Fig. 7. Importantly, some parameter values allow a bi-stable behavior. It is thus possible to understand the direct relationship between the biophysical variations in potassium concentration and the type of bifurcations by observing the trajectory of the membrane potential in this space for periodic events identified previously. During periodic oscillatory behavior, the neuron is initially in resting state (blue plane). The membrane potential slowly increases due to the rise in extracellular potassium, until it reaches a SN (green plane) and then encounters a limit cycle. The slow subsystem then drives it to a negative value of Δ[K]_i_, were the limit cycle meets a SN producing homoclinic bifurcation. These bifurcations are observed at the onset and offset of bursting and SLE behaviors in the model. To have a better understanding of these trajectories, animations with the dynamics of the fast subsystem are available in supplementary material (Fig. S1, S2, S3, S4). We therefore have here a means of bringing together the biophysical aspects, described previously, with the phenomenological vision of dynamical systems approach.

**Table 2.**
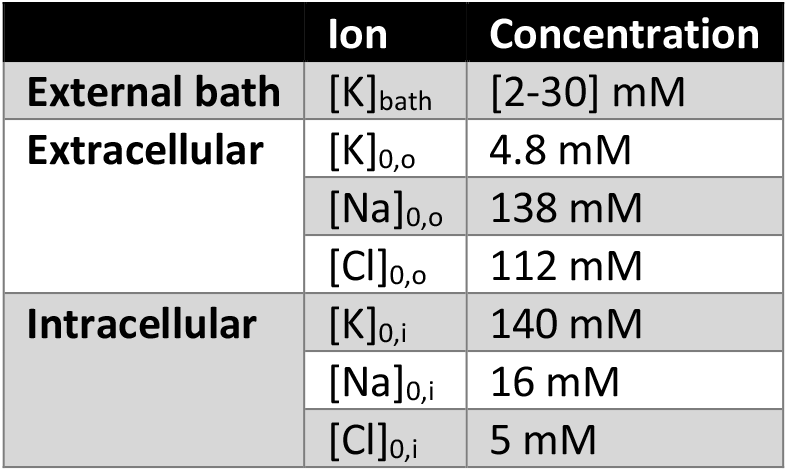
Physiological reference values.

**Figure 7:**
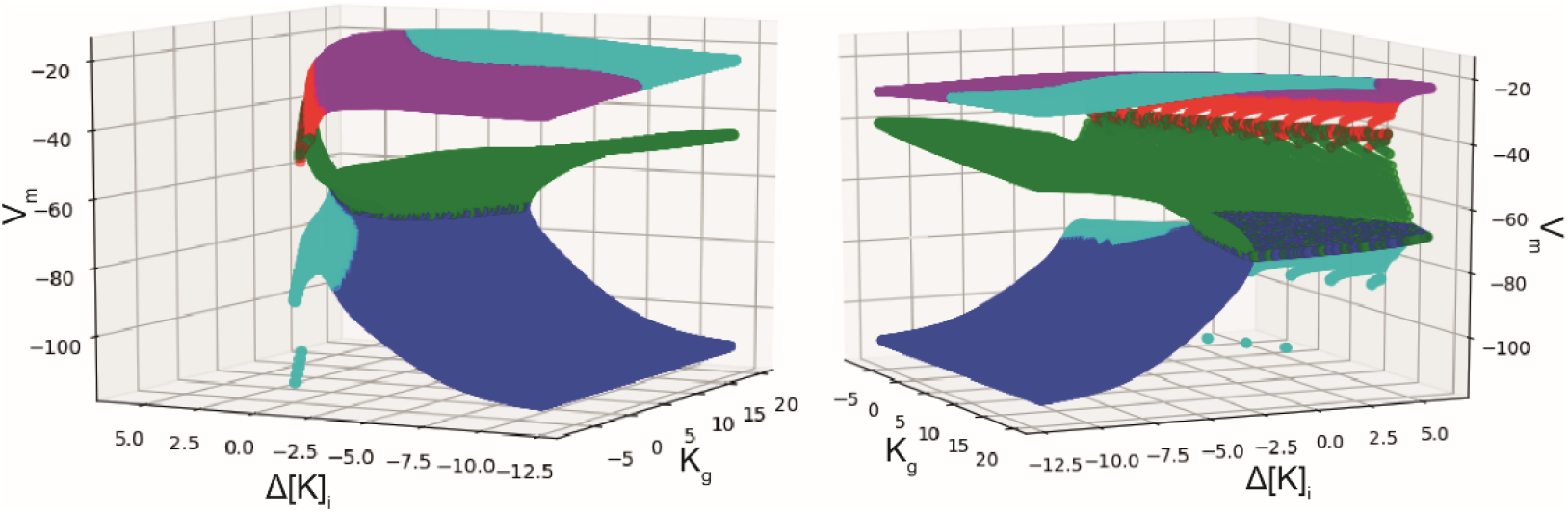
Fixed points of the fast subsystem. Fixed point of the fast subsystem (V_m_) considering the variables from the slow subsystem as parameters. We used a numerical methods with SymPy (Meurer et al. 2017) and SciPy (Millman and Aivazis 2011) libraries, to find the roots and the eigenvalues of the Jacobians of the 2D fast subsystem, and thus the stability considering the existence and the sign of real and imaginary parts of the eigenvalues of the Jacobians. Blue: stable node, green: saddle node, cyan: stable focus, magenta: unstable focus, red: unstable node. Two different angles of view are presented, illustrating the manifold that permits bi-stability.

## Discussion

The aim of this work is to develop a minimal biophysical model at single neuron level based on time scale separation, where the system is able reproduce the dynamics which have been identified in experiments (Bikson et al. 2003; Jirsa et al. 2014; Quilichini et al. 2002; George G. Somjen 2001; Uva et al. 2013) and described by generic models (Jirsa et al. 2014; Saggio et al. 2017). For this purpose, we developed a three-compartment model: a cell equipped with voltage-gated channels to generate action potentials, and Na+/K+ pump to maintain stable ion concentration, an extracellular space surrounding the cell and an external bath that can uptake/release potassium from/to extracellular space. We managed to describe the interaction between these compartments using a system of four differential equations describing two fast and two slow variables. The fast variables delineate excitability while the slow ones, outline potassium changes from the first and third compartments. The sodium concentration changes are not excluded from our model but are linked to potassium through the electroneutrality principle. We have shown that despite its simplicity the model was able to mimic six electrophysiological behaviors classically recorded in neurons and neuronal networks, via the variation of only one parameter. All parameter values were within biophysical ranges (Table1) (Hille 2001; Kandel, Eric R., Schwartz James H., Jessell Thomas M. n.d.). The model has two main limitations. The fast system describes only intrinsic excitability and does not include synaptic currents. And, the slow system is based (only) on potassium concentration. Introducing synaptic inputs would increase the dimension of the system. We propose that synaptic inputs would act as a noise generator increasing the probability to reach the bifurcation as demonstrated experimentally (Jirsa et al. 2014); including them should not change the general behavior of the model. Furthermore, ion homeostasis is not reduced solely to potassium. Potassium is just one candidate among many others for the slow system. Numerous studies have reported large changes in concentration of Ca^2+^ (Heinemann et al. 1986), Cl^-^ (Miles et al. 2012; Raimondo et al. 2015) and neurotransmitters during seizures (Chapman et al. 1984; During and Spencer 1993). Likewise, decreasing extracellular Ca^2+^ leads to seizures (Jefferys and Haas 1982), which are characterized by SN/homoclinic bifurcations (Jirsa et al. 2014). Since it is possible to trigger similar SLEs via totally different biophysical mechanisms (Jirsa et al. 2014), we propose that the K^+^-dependent mechanism we describe, is one among many the possible paths leading to the same end point. In our model, changes in potassium constitute the causal factor driving the neuron through different types of activities. Although similar changes in potassium are measured experimentally when networks (and not cells) undergo such transitions, causality has not been demonstrated experimentally, only correlation. Another limitation exists due to the formalism used. If [K]_bath_ tends to zero then membrane potential hyperpolarize until the Nernst potential are no longer defined due to division by zero. We reach here the limit of the conductance-based model from Hodgkin-Huxley formalism. Due to the expression of the Nernst potential, if the ratio [K]_o_/[K]_i_ approaches zero, then the I_K_ current increases towards infinity, which is not physiologically plausible. Another factor to consider is that the dynamics of the single cell is driven by slow changes of extracellular variables, which, in a biological system, is shared with neighboring cells. So, these slow variables can also be responsible for the genesis of network activity (Naze et al. 2015). As these mechanisms exist both at the network and single neuron level, it would be simplistic to conclude that a seizure at the network level is due to the combined expression of seizures at the single cell level. Since a neuronal network can be seen as a complex system of many components, coupled in a non-linear manner, seizures may just be an emergent property, perhaps taking advantage of the fact that they are already encoded at the single cell level. The same consideration applies to other pathological activities such as SIA and DB, which corresponding pattern have been found in dynamics of our model.

However, we only studied the dynamics for variations of few chosen parameters based on physiological observations identified in previous works. The parameters explored here show that the model can produce different combinations of onset/offset bifurcations. Numerous studies used ion concentration variations in biophysical models to generate various types of activity (Barreto and Cressman 2011; Bernard et al. 2014; Cressman et al. 2009; Florence et al. 2009; Krishnan et al. 2015; Øyehaug et al. 2012; Yina Wei, Ullah, and Schiff 2014; Yina Wei, Ullah, Ingram, et al. 2014). Descriptions of ion concentration dynamics for bursting have been done by Barreto *et al*. (Barreto and Cressman 2011), based on a slow/fast system. In this work, the bifurcations for SLEs are SNIC and Hopf. This approach, based on ion concentration dynamics, permits the unification of spike, seizure and spreading depression proposed by Wei and al. (Y Wei et al. 2014). As different models can lead to similar dynamics (Prinz et al. 2004), this may suggest that different minimalist models are possible to obtain a unified framework. In our work, we proposed a conductance-based model of the neuronal membrane, exhibiting an extended repertoire of behavior and introducing sustained ictal activity in a unified framework. Another difference with previous work is that our model can exhibit bi-stable modes saddle-node/homoclinic bifurcations, which are the most commonly observed in recordings from patients and experimental animal models (Jirsa et al. 2014). Our model does not take into account variation of volume or oxygen homeostasis as in (Y Wei et al. 2014) but, only variations of ion concentrations, driven by diffusion of potassium from EB. It seems intuitive that other biological variables could be considered as slow variables to drive the fast subsystem in a reduced biophysical model. The work of Øyehaug*et al*.(Øyehaug et al. 2012) presents interesting dynamical features with saddle-node/homoclinic bifurcations for SLEs. However, this model is much more complex as it describes numerous biological features and mechanisms. In comparison to previous works (Barreto and Cressman 2011; Krishnan et al. 2015; Øyehaug et al. 2012; Yina Wei, Ullah, Ingram, et al. 2014), our model is reduced to only four equations. We sought to include only a minimal number of mechanisms necessary to reproduce neural dynamics. Chizhov *et al*. (Chizhov et al. 2018) proposed a biophysical model (Epileptor-2) of ictal activities based on the Epileptor (Jirsa et al. 2014), using different differential equations. In high potassium conditions, Epileptor-2 produces bursts of bursts, described as ictal-like discharges. However, the most common form of seizure belongs to the saddle-node/homoclinic form, which starts with low voltage fast activity, and ends with bursts slowing down in a logarithmic fashion. The latter was reproduced in the present model, including the period during which neurons stop firing (depolarization block) after seizure onset. Another difference lies in K^+^ dynamics. In Epileptor-2, neuronal firing ends when extracellular K^+^ returns to baseline level (see Fig 10. in (Chizhov et al. 2018)), whereas in the present model, there is a delay, as consistently found experimentally, as a result of glial cell action. This phenomenon in our model can be visualized by observing the evolution of [K]_o_ in Fig. 4. Although the Epileptor-2 is not an “intrinsic” Slow/Fast dynamical system, indeed, this model does not describe an independent node as it takes in account the external influence from synaptic inputs from neuronal population. In our model, the observed dynamics, is only due to internal interactions between three compartments.

In conclusion, we developed a biophysical model of a single neuron that, despite its simplicity, is able to generate, in a unified framework, many patterns of neuronal network activity found in experimental recording as well as in generic mathematical models. We show that transition from physiological to paroxysmal activity can be obtained by variation of model parameters relating to ion homeostasis while excitability parameters remained constant. Thus, we proposed a simple biophysical model comparable to generic models (El Houssaini et al. 2015; Jirsa et al. 2014; Saggio et al. 2017), offering the possibility of a biological interpretation of observed dynamics. Neuronal networks increase in complexity from flies to humans, but the basic properties of neurons are roughly conserved. The present study shows that acting on an external variable allows single neurons to go through various patterns of activities, which are also found at the network level in the form of seizures, sustained ictal activity and depolarization block (Cunliffe et al. 2015; Jirsa et al. 2014). We propose that they constitute one of the most primitive forms of activities, appearing as soon as neurons are present.

## Materials and Methods

In this project we aim to build a minimal biophysical model that describes different electrophysiological states of a single neuron, the model is schematized in Fig.1. The model describes three compartments: the intracellular space (ICS), the extracellular (ECS) space and the external bath (EB). Parameters chosen correspond to values observed in whole cell recording. The ion exchange between the ICS and the ECS is carried out by the current flowing through the sodium, potassium, and chloride voltage-gated channels (eq.(5),(6) and (7)), and by the sodium-potassium pump generated current (eq.(8)). Parameters values for these currents have identified in (Hamada et al. 2003; Hille 2001; Läuger 1991) and the membrane capacitance in (Golowasch et al. 2009). Passive diffusion of potassium exists (eq.(4)), between EB and ECS. The EB is mimicking the K+ buffering of vasculature/astrocytes. In ICS and ECS actualization of potassium and sodium concentrations are done (eq.(14)-(20)). The γ parameter has the same unit as the inverse of the Faraday constant, and it is a scaling parameter that permit to include all the mechanisms not detailed in this model which affect the concentration variations (such as co-transporter, exchangers). The values of all the parameters used are given in table 1 and physiological reference and initial values are given in table 2 and table 3.

**Table 3.**
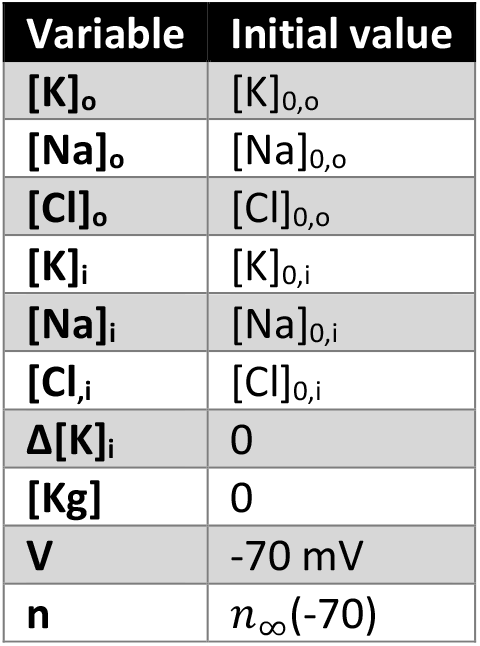
Initial values.

The model is a slow-fast dynamical system based on 4 equations. The fast system describes the membrane potential eq.(1) and potassium conductance gating variable eq.(2). The slow system describes intracellular potassium concentration variation eq.(3) and extracellular potassium buffering by external bath eq.(4).

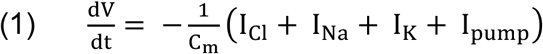

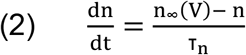

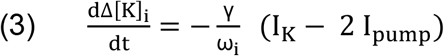

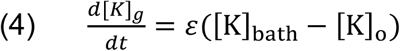

With currents:

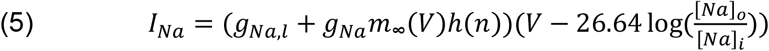

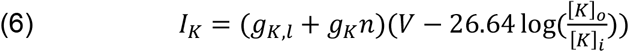

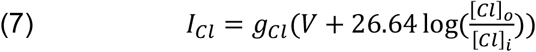

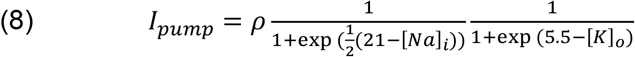

And conductance variables:

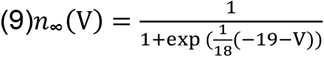

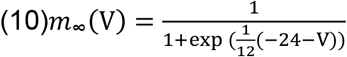

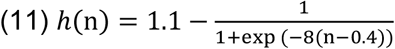

The fast subsystem of the model, (eq. (1)&(2)), is a reduction and simplification of conductance-based models, first describe by Hodgkin–Huxley (HH). From the original publication (Hodgkin, A. L., Huxley 1952) the activation variable of K+ channels is determined by the equation (eq.12):

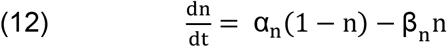

Where β(V) and α(V) are the voltage-dependent rate constants determining the probability of transitions between, respectively, opened and closed state of the ion channel. To simplify the model, we propose to describe the variable n, through the voltage-dependent parameter n_inf_(V) and a constant parameter τ_n_. In our model, n_inf_(V) is the probability to find a channel at open state at a given membrane potential while τ_n_ is the fixed time constant that described the speed for channels to respond to the change of membrane potential. Based on available data in the literature (Bekkers 2000; Hodgkin, A. L., Huxley 1952), and considering that the mean number of channels opened at a given potential is constant, we could qualitatively estimate this relationship (eq.9). In the HH model, the time constant is dependent on the membrane potential due to the formalism used (eq.12). The HH model has been build thanks to experiments done on the squid giant axon, which present differences from on recording of mammalians neurons. We compare the n_inf_(V) of our model and 1/τ(V), and n_inf_(V) of the HH model in Fig. 8(a). The shape has been kept from the HH model but starts to increase for lower values of membrane potential. For the voltage-gated sodium channels, variables for opening, m, and for closing, h, have been described(Hodgkin, A. L., Huxley 1952). With the same logic, we can consider the percentage of all population of channels opened. But because this is a very fast mechanism (Hille 2001), it can be considered as an instantaneous function of V (E. Izhikevich 2007) (eq.10). Krinskii and Kokoz (Krinskii,V.I., Kokoz 1973) showed that n(t)+h(t) is almost constant, so h can be considered as a function of n. Because of the previous modification, we adapted this fitting to obtain the equation of h(n) (eq.11). Due to these simplifications, the interdependence of gating variables makes the spiking rate dependent on τ, as shown in Fig.8(b).

**Figure 8:**
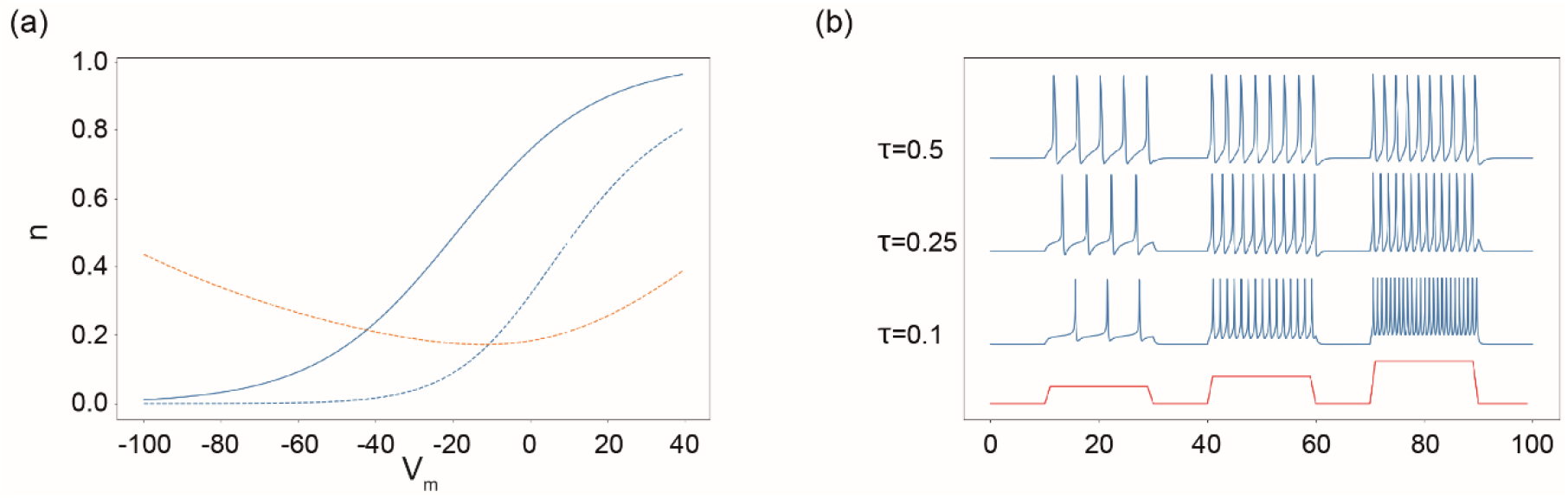
modification in gating variables. (a) n_inf_ of our model in blue, and n_inf_ and 1/τ of the Hodgkin-Huxley model respectively in dash blue and red, function of the membrane potential. (b) Response of the fast subsystem of our model to step current stimulation (red) with three different values of τ (0.1, 0.25, 0.5 ms). The value of τ influence the frequency rate spike for a same injected current.

To be able to take into account concentration variation limiting the number of equations we applied reductions. Inspired by the work of Hübel (Hübel 2015; Hübel and Dahlem 2014), electroneutrality permits the Eq.(13), and so to the Eq.(14). The ratio (C_m_ γ)/ω_i_ is very small (<10^−5^) and so, the right-hand side of Eq.(14) could be considered to be zero. The chloride concentration changes are assumed to be small and regulated by mechanisms which are not described in our model (Doyon et al. 2016). So, in our model, the chloride concentration remains constant.

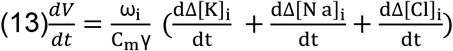

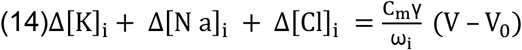

Thanks to these reductions, concentration variations are calculated as follow:

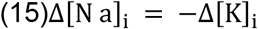

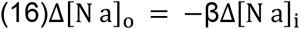

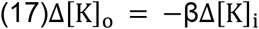

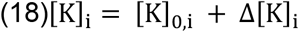

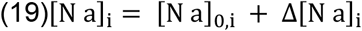

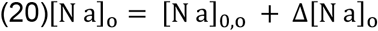

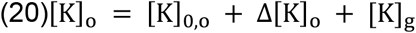

## Acknowledgement

This research was partially supported by the European Union’s Horizon 2020 research and innovation program under grant agreement No. 785907 (SGA2), and No. 945539 (SGA3) Human Brain Project and was also financed in part by the “Coordenação de Aperfeiçoamento de Pessoal de Nível Superior - Brasil (CAPES) - Finance Code 001”, the National Reasearch Council “Conselho Nacional de Desenvolvimento Científico e Tecnológico (CNPq)” and the Family Allowance Fund (CAF de Marseille, France).

The authors declare no conflict of interest.

## Supporting information

**S1 Animation. Dynamics of the membrane potential during burst**. Considering the two slow variables as parameters of the fast subsystem, fixed point has been found: blue: stable node, green: saddle node, cyan: stable focus, magenta: unstable focus, red: unstable node. The system starts at a stable fixed point and is slowly driven to cross a saddle-node and then follow a limit cycle, until it cross again a saddle-node (creating the Homoclinic bifurcation), and go back to a stable fixed point.

**S2 Animation. Dynamic during Burst observed in the phase plane**. The n nullcline (blue line) and the V nullcline (blue points) solved numerically. The system starts at a stable fixed point and is slowly driven to cross a saddle-node and then follow a limit cycle, until it cross again a saddle-node (creating the Homoclinic bifurcation), and go back to a stable fixed point.

**S3 Animation. Dynamics of the membrane potential during SLEs**. Considering the two slow variables as parameters of the fast subsystem, fixed point has been found: blue: stable node, green: saddle node, cyan: stable focus, magenta: unstable focus, red: unstable node. The system starts at a stable fixed point and is slowly driven to cross a saddle-node and then follow a limit cycle, it cross successively two Hopf bifurcations to come back to a limit cycle until it cross again a saddle-node (creating the Homoclinic bifurcation), and go back to a stable fixed point.

**S4 Animation. Dynamic during SLEs observed in the phase plane**. The n nullcline (blue line) and the V nullcline (blue points) solved numerically the system start at a stable fixed point and is slowly driven to cross a saddle-node and then follow a limit cycle, it cross successively two Hopf bifurcations to come back to a limit cycle until it cross again a saddle-node (creating the Homoclinic bifurcation), and go back to a stable fixed point.

