## Supplementary figures and images for "A unified physiological framework of transitions between seizures, sustained ictal activity and depolarization block at the single neuron level"

### S1

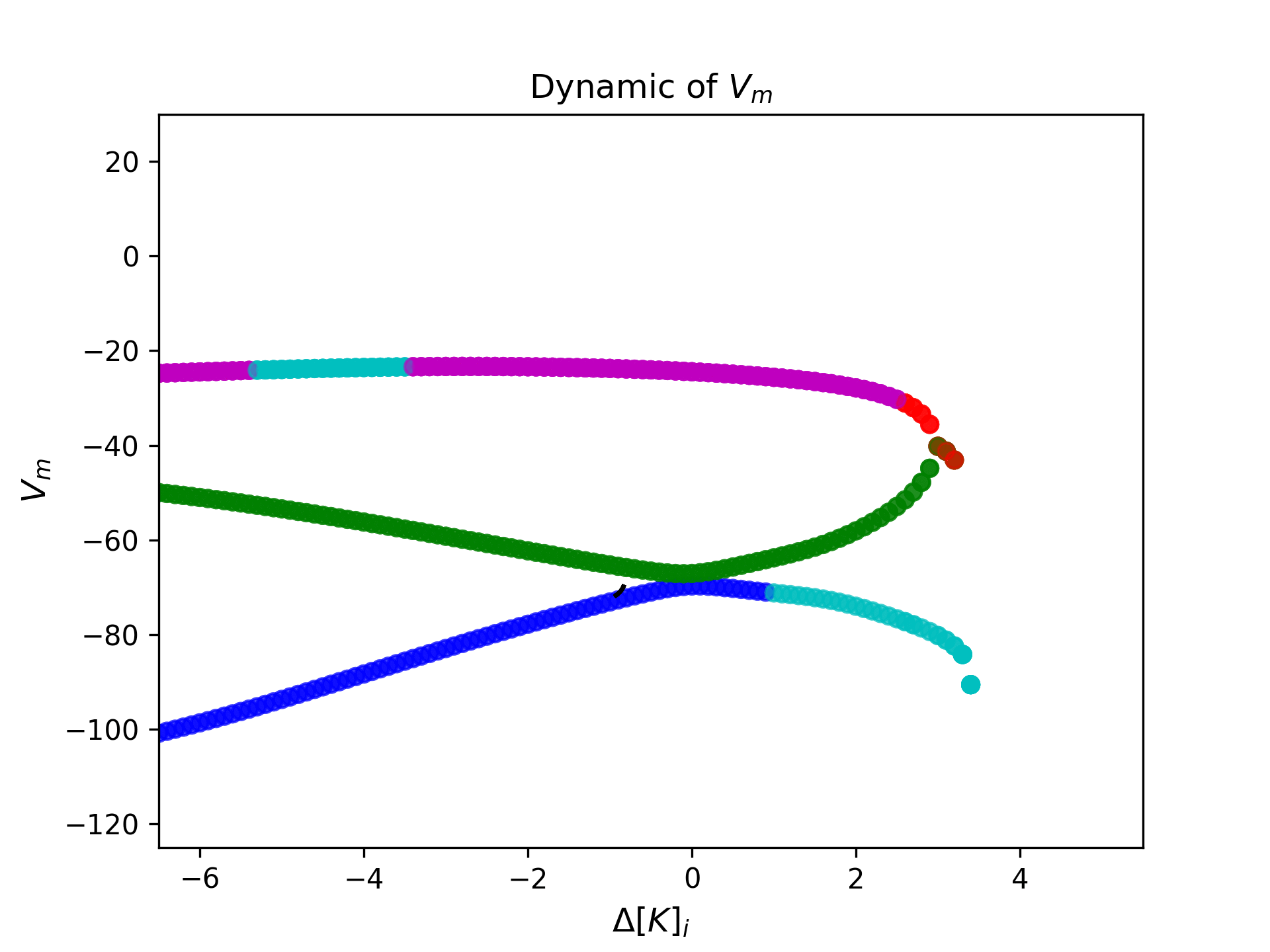

### S2

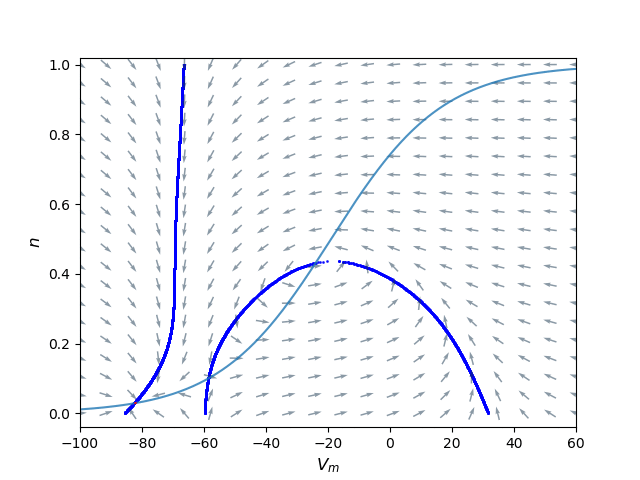

### S3

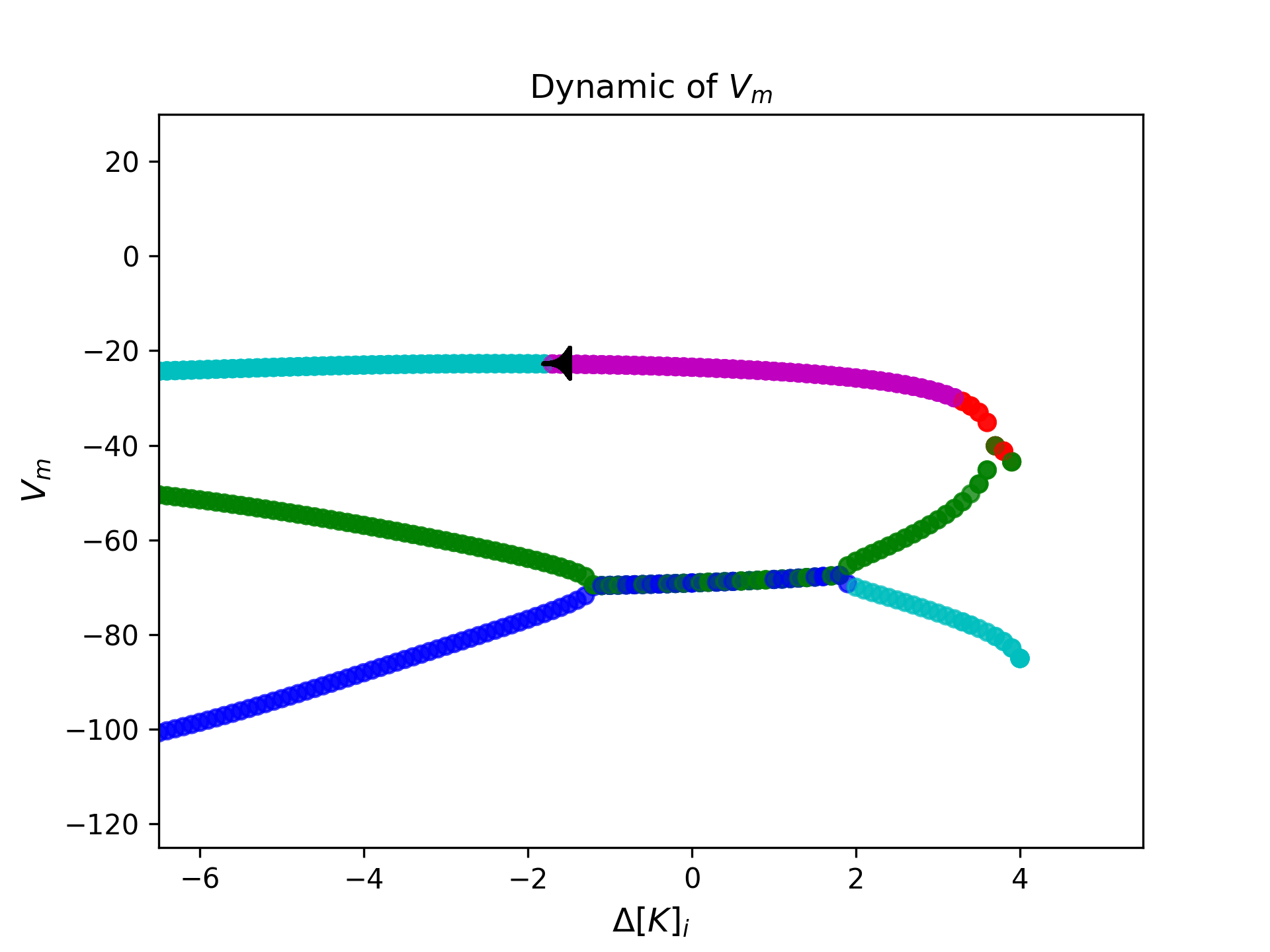

### S4

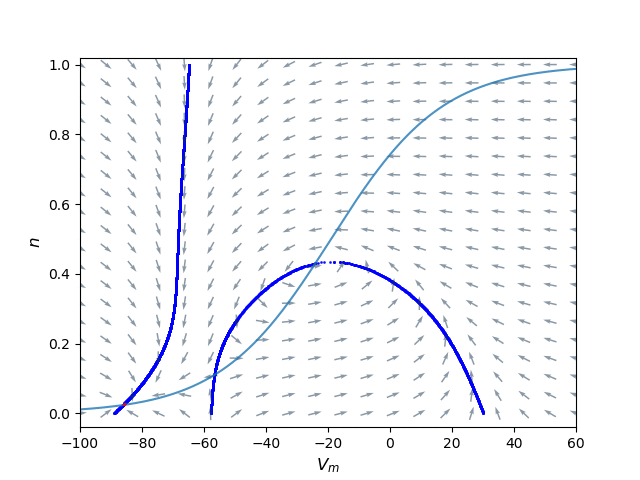
